# Heterogeneous Appetite Patterns in Depression: Computational Modeling of Nutritional Interoception, Reward Processing, and Decision-Making

**DOI:** 10.1101/2024.09.25.614954

**Authors:** Yuuki Uchida, Takatoshi Hikida, Manabu Honda, Yuichi Yamashita

**Affiliations:** Department of Information Medicine, National Institute of Neuroscience, National Center of Neurology and Psychiatry, Tokyo, Japan; Medical and Dental Sciences, Graduate School of Medical and Dental Sciences, Tokyo Medical and Dental University (TMDU), Tokyo, Japan; Laboratory for Advanced Brain Functions, Institute for Protein Research, Osaka University, Osaka, Japan

**Author notes:** **Correspondence:** Yuichi Yamashita.

**Keywords:** appetite, computational neuroscience, computational psychiatry, decision-making, dopamine, homeostasis, homeostatic reinforcement learning

## Abstract

Accurate interoceptive processing in decision-making is essential to maintain homeostasis and overall health. Disruptions in this process have been associated with various psychiatric conditions, including depression. Recent studies have focused on nutrient homeostatic dysregulation in depression for effective subtype classification and treatment. Neurophysiological studies have associated changes in appetite in depression with altered activation of the mesolimbic dopamine system and interoceptive regions, such as the insular cortex, suggesting that disruptions in reward processing and interoception drive changes in nutrient homeostasis and appetite. This study aimed to explore the potential of computational psychiatry in addressing these issues. Using a homeostatic reinforcement learning model formalizing the link between internal states and behavioral control, we investigated the mechanisms by which altered interoception affects homeostatic behavior and reward system activity via simulation experiments. Simulations of altered interoception demonstrated behaviors similar to those of depression subtypes, such as appetite dysregulation. Specifically, reduced interoception led to decreased reward system activity and increased punishment, mirroring the neuroimaging study findings of decreased appetite in depression. Conversely, increased interoception was associated with heightened reward activity and impaired goal-directed behavior, reflecting an increased appetite. Furthermore, effects of interoception manipulation were compared with traditional reinforcement learning parameters (e.g., inverse temperature *β* and delay discount *γ*), which represent cognitive-behavioral features of depression. The results suggest that disruptions in these parameters contribute to depressive symptoms by affecting the underlying homeostatic regulation. Overall, this study findings emphasize the importance of integrating interoception and homeostasis into decision-making frameworks to enhance subtype classification and facilitate the development of effective therapeutic strategies.

## 1 Introduction

Interoception that is appropriately integrated into decision-making is essential for maintaining homeostasis and overall health (Cannon, 1929; Friston, 2013; Stephan et al., 2016). Maladaptive homeostasis is associated with eating disorders (Brown et al., 2017; Khalsa et al., 2022), unbalanced feeding in autism spectrum disorder (Fiene and Brownlow, 2015), and depression (Paulus and Stein, 2010; Avery et al., 2014; Stephan et al., 2016). Among these conditions, nutritional homeostasis dysregulation could be a primary diagnostic marker of depression, which is characterized by symptoms of maladaptive appetite and is used as a criterion for classification (Weissenburger et al., 1986; Zimmerman et al., 2011; American Psychiatric Association, 2022). Maladaptive appetite in depression is heterogeneous, either increasing or decreasing in different cases (Maxwell and Cole, 2009; Simmons et al., 2013, 2016, 2020).

Nutrient homeostatic dysregulation underlying depression has been actively studied in recent years as it is crucial for effective subtype classification of depression and development of appropriate treatments (Konttinen et al., 2010; Privitera et al., 2013; Cosgrove et al., 2020). For example, from a neurophysiological perspective, research has revealed that changes in appetite in individuals with depression are related to altered activation of the mesolimbic dopamine system and areas that are strongly associated with interoceptive processing, such as the insular cortex (Simmons et al., 2016, 2020). These findings suggest that alterations in reward processing and the complex interplay between nutrient interoceptive processing are involved in changes in nutrient homeostasis and altered appetite in depression. However, system-level mechanisms, including those affecting brain activity and behavior, remain largely unclear (Young et al., 2021).

To address these challenges a neurocomputational theory-based methods to reveal the pathophysiology of neuropsychiatric conditions (i.e. “computational psychiatry”) is expected to make important contributions (Montague et al., 2012; Friston et al., 2014; Yamashita, 2021; Takahashi et al., 2023). In the field of depression research, there has been growing interest in using the reinforcement learning (RL) theory in combination with behavioral experiments to test hypotheses and further our understanding of the underlying mechanisms (Takahashi et al., 2008; Kunisato et al., 2012; Toyama et al., 2019). However, the integration of interoception and homeostasis into the theoretical frameworks of decision-making has not progressed sufficiently (Paulus, 2007; Rangel, 2013; Morville et al., 2018).

Therefore, in this study, we aimed to provide a systems-level explanation for the dysregulation of nutrient homeostasis in depression using computational psychiatry methods to clarify the mechanisms by which changes in interoceptive processing and alterations in reward system activity are related. Specifically, we attempted to integrate the mechanisms of homeostasis and decision-making and provide a system-level explanation of the functions of the reward system and nutritional state. To achieve this, we used the homeostatic RL (HRL) model, which formalizes the relationship between internal states and the drive that controls behavior as the “homeostatic space” (Keramati and Gutkin, 2014; Keramati et al., 2017; Morville et al., 2018; Hulme et al., 2019; Uchida et al., 2022). Using the HRL model, we aimed to interpret the changes in homeostatic maintenance behaviors and reward system activity related to changes in interoceptive sensations by conducting a pseudomanipulation experiment (simulation) of reduced and exaggerated interoceptive sensations in the HRL model. In addition, we aimed to compare the effects of the modulation of RL parameters previously associated with depression on decision-making behaviors, focusing on the effects of changes in interoceptive processing within the HRL model. This comparison allowed us to understand the bias in decisionmaking behavior caused by RL parameters in relation to changes in the behavior of the model resulting from alterations in interoceptive sensation. Through this study, we hope to offer a systems-level explanation of the associations among phenomena previously considered as changes in RL parameters, changes in interoceptive processing, and nutrient homeostasis in depression.

## 2 Materials and Methods

### 2.1 HRL Model

In this study, nutritional homeostasis was modeled using the HRL model. This model assumes that homeostasis is an RL process, in which the minimization of deviations in internal states from an optimal level (i.e., homeostasis) is treated as a computation for maximizing the sum of rewards. In the HRL model, a multidimensional metric space in which each dimension represents an internal state (such as body temperature, blood glucose density, water balance, and sodium level) is defined as a “homeostatic space”. In this homeostatic space, the drive function *D(H*_*t*_) is defined as the distance between the internal state of the *i*-th component (e.g., water or sodium) at time *t*, 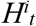, and the ideal internal state *H*^*\*i*^:

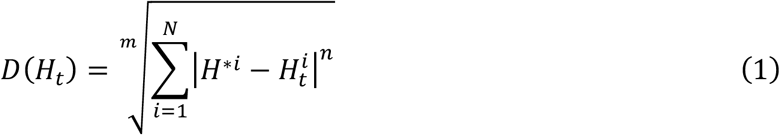

where *m* and *n* are free parameters that define the distance and *N* is the total number of dimensions for the internal states (e.g., water and sodium). When the internal state approaches the ideal state, the drive function decreases. Based on this drive function, the reward *r*_*t*_ is determined as the change in the values of the drive function from time *t* to time *t+1*. Specifically, to implement nutrient intake, the internal state at time *t+1* should contain the amount of nutrient intake at time *t*, defined as *K*_*t*_.

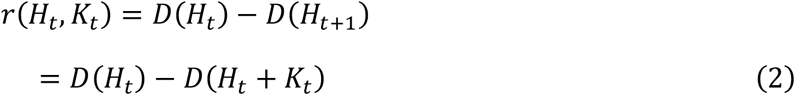

As described later, in the HRL model, the intake of taste stimuli 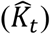 can be modeled as a predictor of the actual nutrient intake (*K*_*t*_). Under this assumption, the reward is calculated as follows:

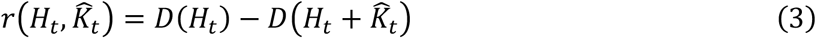

Q-learning was used to model the RL process. In this model, the values of action *a*_*t*_ (e.g., intake, do nothing…) and *Q*_*t*_*(a*_*t*_) are updated based on the temporal difference error (*δ*_*t*_):

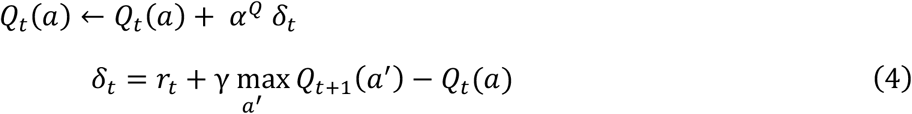

where *α*^*Q*^ is the learning rate for *Q*_*t*_*(a), a’* is the next candidate action, *δ*_*t*_ is the TD error, and γ is the discount rate. Action selection depends on the relative magnitudes of the values of each action (*Q*-value) according to the softmax function.

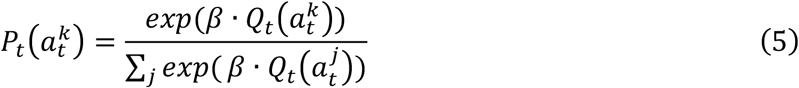

where *P*_*t*_ *(a*^*k*^) is the probability of an action *a*^*k*^ to be selected at time *t*, and *β* is the inverse temperature, a parameter controlling the randomness of an action. The initial values of the *Q*-values of both actions were set to *0*. Therefore, the first action is chosen at random. When the agent intakes, the internal state increases with *K*_*t*_, which is a constant that defines the amount of intake. When nothing was chosen, *K*_*t*_ was set to *0*. At *t = 0*, representing the hungry state, the first internal state (*H*_*0*_ *= 100*) was far lower than the ideal state (*H*^*\**^ *= 200*). At this stage, the drive function is large because it corresponds to the distance from the internal state at time *t* (*H*_*t*_) to the ideal state (*H*^*\**^ *= 200*; Eq. 1). If an agent performs the intake behavior at this moment, the internal state is expected to increase and the drive function is anticipated to decrease, resulting in a positive reward (Eq. 3). In addition, the natural decrease in nutritional balance was implemented as follows using the temporal decay constant *τ* (Eq. 6):

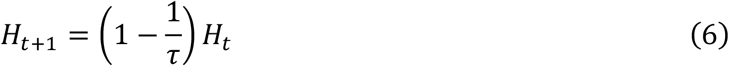

The reward value was calculated as follows (Eq. 7):

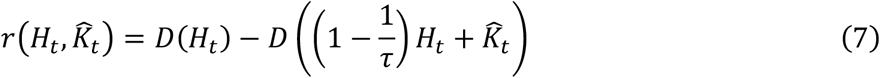

After updating the *Q*-values via Q-learning, the agent selects the next action. As previously mentioned, the HRL model assumes that 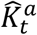, the cognition of the stimulus based on the reward from the action *a*_*t*_, is renewed through learning. In this study, the following equation was used to update 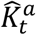:

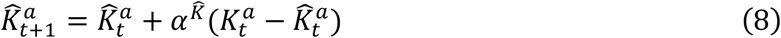

The detailed values of the simulation parameters are listed in **Supplementary Table 1**.

### 2.2 Nutritional Homeostasis: Intake-After-Food-Restriction Task

We performed an intake-after-food-restriction task to investigate the applicability of the HRL model to nutritional homeostasis. The computational algorithm is illustrated in Figure 1B. For simplicity, only one nutritional state is considered. The external state (*S*_*0*_) and two actions–do nothing (*a*^*0*^) and intake (*a*^*1*^)–were assessed (Figure 1A).

**Figure 1:**
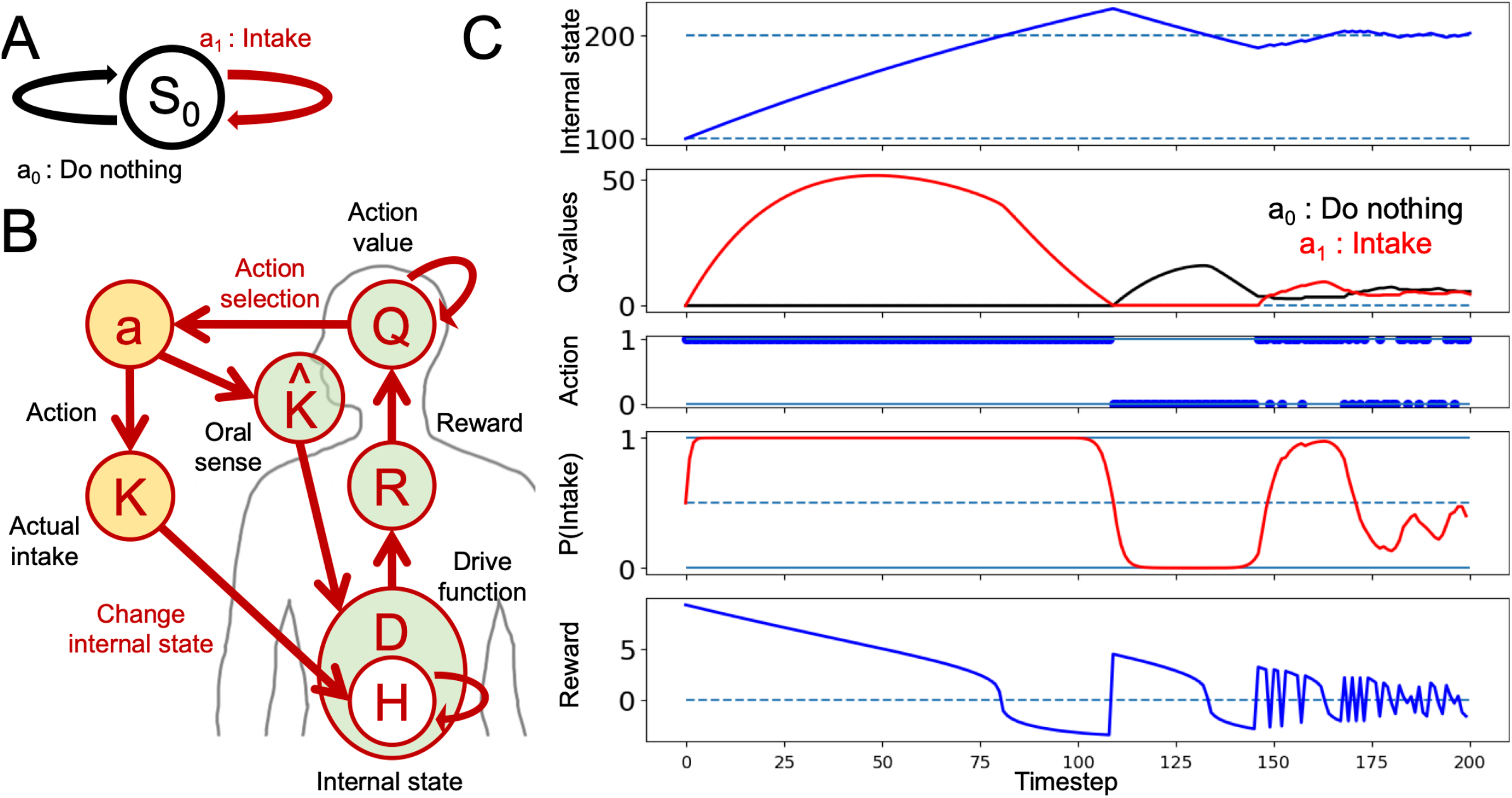
Nutritional homeostatic maintenance according to the homeostatic reinforcement learning (HRL) model. **(A)** Definition of the state and two actions in intake-after-food-restriction simulations. **(B)** Schematic of the computational process of the HRL model. **(C)** Example of homeostatic behavior. Changes in the internal nutritional state (*H*), value of each action (*Q*-value), selected actions (*a*), probability of intake (*P(Intake)*), and magnitude of reward (*R*) are plotted. Solid lines indicate the results of a single trial. Dotted lines in the panel of the internal state indicate the ideal point (*H*^*\**^ *= 200*) of the nutrient. In the panel related to actions (*a*), action *1* indicates “intake”, and action *0* indicates “do nothing”. At the beginning of the simulation, internal nutritional state was *100*, and *Q*-values for each action were set to *0*. After several random selections of action, *Q*-value of nutrient intake was increased, and the internal nutritional state quickly reached the ideal point, maintaining homeostatic regulation of behavior.

The following formula was adopted to implement the alteration in interoception:

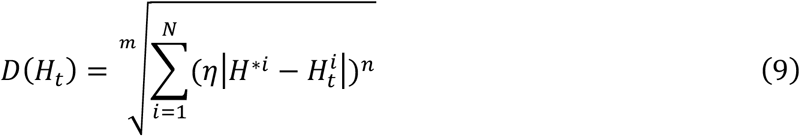

where *η* is a parameter that refers to the modulation of the interoception and is a constant over the difference between the ideal and actual internal state. The validity of this implementation is discussed in the Discussion section.

### 2.3 Mountain-Climbing Task

Mountain tasks have been utilized to assess whether individuals prioritize short-term, small rewards or long-term, large rewards that can only be obtained by enduring sequences of punishments (e.g., “Mountain car”) (Sutton and Barto, 2018). In this study, a derived form was employed for this task. The task comprised *8* states (*S*_*0*_*-S*_*7*_; Figure 3A). For each condition (control and low interoception), the experiment consisted of *30* trials, and each trial included *15* episodes (Figure 3B).

At the beginning of each episode in this task, the agents started at *S*_*0*_ with an internal state of *100* (*H*_*0*_ *= 100*). The agent can only choose to intake at two states: the bottom state (*S*_*0*_), where it can choose to consume a small amount, and the summit state (*S*_*7*_), where it can choose to consume a large amount. At *S*_*0*_, the agents have two options for selecting an action: *a*^*00*^ (small intake) or *a*^*01*^ (moving horizontally). In *S*_*1*_-*S*_*6*_, the agents have the option to climb or descend (and move horizontally only in *S*_*1*_). It is important to note that climb actions (*a*^*10*^, *a*^*20*^, …, *a*^*60*^) result in a constant decrease in the internal state, which acts as a punishment in the context of nutritional deficiency accompanied by climb actions. Other actions, than the climbs, follow decrease in internal state derived from only the attenuating rate (*τ*), but the decreases with climb actions resulted from both *τ* and constant cost additionally. Once the agent selects *a*^*70*^ (large intake), the episode ends, the state-action values (*Q*) and the prediction of the internal state increase (*K*^*^*^) are carried over to the next “episode”. The initial states of the subsequent episodes in the mountain-climbing task were consistently set to *S*_*0*_, and the episodes were iterated 15 times. One trial of the mountain-climbing task was completed when *15* large-intake actions (*a*^*70*^) were performed (Figure 3A, B). Each condition consisted of *30* trials (Figure 3B). A different set of free parameters was used in the mountain-climbing task than in the intake-after-food-restriction task to ensure that the number of time steps in a trial converged within rational time steps (Supplementary Tables 1 and 2).

## 3 Results

### 3.1 Nutritional Homeostasis: Intake-After-Food-Restriction Task

First, we demonstrated the behavior of a healthy control model using the intake-after-food restriction task (Figure 1). The detailed process is described in the **Materials and Methods** section.

At the beginning of the simulation, the internal nutritional state was set to a value far from the ideal state (corresponding to a fasting state), and *Q*-values for each action were set to *0*. After several random selections of actions, *Q*-value of nutrient intake increased, and internal nutritional state quickly reached the ideal point, maintaining the homeostatic regulation of behavior. The value of do nothing decreased, and *Q*-values of the intake increased and remained relatively high for some time, even after exceeding the set point (*H*_*t*_ *> H* ^*\**^ = *200*). Subsequently, frequency of doing nothing increased owing to continued punishment from an excess internal state as the value of doing nothing became greater than the *Q*-value of intake. Continued do nothing caused the internal state to decline due to natural decay. In response to this decline, *Q*-value of intake was greater than that of doing nothing, resulting in the maintenance of homeostatic regulation (Figure 1C). This simulation expressed one aspect of nutritional homeostatic maintenance behavior: the frequency of intake increased after experiencing nutritional deficiency and decreased after achieving sufficiency.

### 3.2 Altered Interoception in the Intake-After-Food-Restriction Task

Next, we investigated the impact of altered interoception on feeding behavior and nutritional homeostasis using an intake-after-food-restriction task. In the simulation, we assessed the simple behavioral characteristics of each model under nutrient deprivation at *100* time points (Figure 2) and quantified the changes in feeding behavior and nutritional homeostasis induced by altered interoception. As shown in Eq. 9, altered interoception was simulated by varying the value of parameter *η* (*η*: : 0 < *η* < 2.0), which is a constant parameter manipulating the impact of the difference between the setpoint (*H*^*\**^ = 200) and actual internal state.

**Figure 2:**
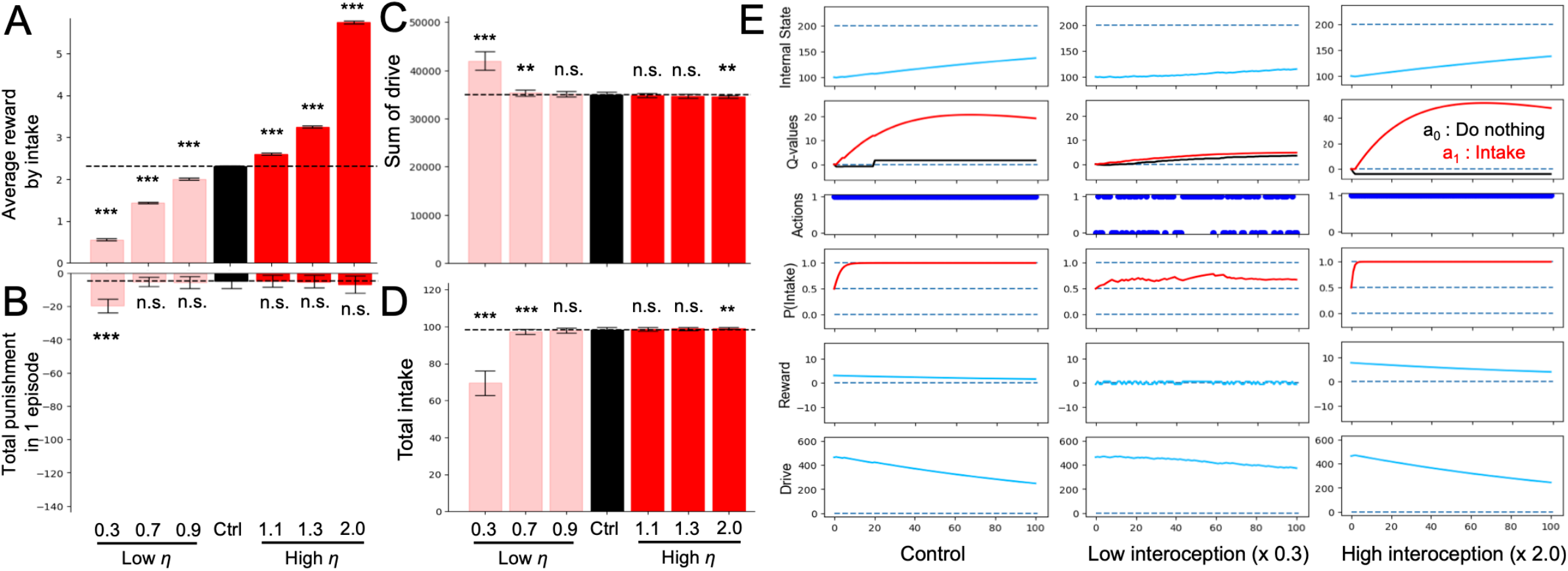
Altered interoception in the intake-after-food-restriction task. **(A)** Average of reward with all intake behavior in a single episode determined from the simulations with altered interoception models. **(B)** Sum of punishments during one episode with altered interoception. **(C)** Sum of drive in one episode determined from the simulated lesion models with altered interoception. **(D)** Total intake in an episode with altered interoception. **(E)** Example of transition of variable from episodes of the control, low interoception, and high interoception models. In panels (A–D), Student’s *t*-test or Welch’s *t*-test was used for between-group comparisons after Levene’s test. \*\**P* < 0.01 and \*\*\**P* < 0.001 (N = 40).

**Figure 3:**
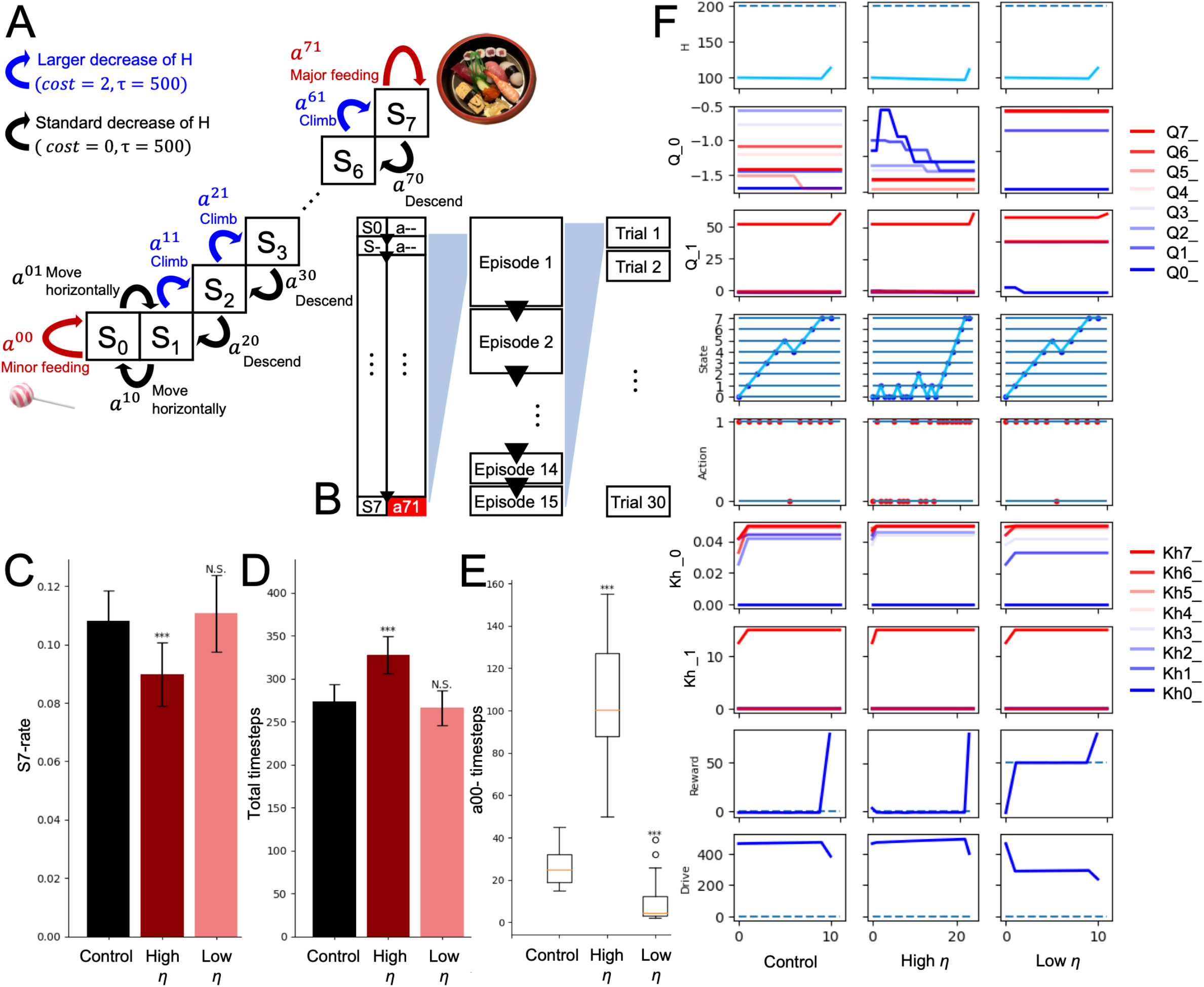
Altered interoception in the mountain-climbing task. **(A)** Definitions of eight states and two actions at each state in the mountain-climbing task. **(B)** Time series relationships among variables. **(C)** S7-rate of the control, low interoception, and high interoception models. **(D)** Total timesteps per trial. **(E)** Total number of minor intakes. **(F)** Trajectories of each variable in the *5*th episode of the control, low interoception, and high interoception models. Significance was determined using the Student’s *t*-test (B, C) or Wilcoxon rank-sum test (D) (30 trials). \*\*\**P* < 0.001; *N*.*S*., not significant.

Figure 2A demonstrates the reward properties showing the average reward per intake (i.e., total reward obtained for an intake within an episode divided by the number of intakes). Figure 2B shows the sum of the punishments in an episode. In the low *η* model (*η* < 1.0), average reward per intake was decreased (Figure 2A), and the sum of punishment was increased (Figure 2B). This occurred because the absolute value of the drive (*D*) obtained by the behavior was smaller (Figure 2E), and consequently, the absolute value of reward (*R*) (i.e., the difference in drive (*D*)), also became smaller (Figure 2E). In addition, the low *η* models reduced the frequency of the optimal action selection (i.e., intake in the case of nutritional deficiency; Figure 2D) because the input of nutritional deficiencies into the model was reduced. In the “Actions” of Figure 2E, action *a*_*1*_ represents intake, and it can be observed that the frequency of intakes decreased in the low interoception (*η* = 0.3) model. Consistent with the decreased intake frequency, the total number of drivers per episode increased (Figure 2C).

In contrast to the low *η* models (*η* < 1.0), the high *η* model (*η* > 1.0) exhibited an increase in the average reward per intake (Figure 2A) and a decrease in the sum of punishment (Figure 2B), reflecting opposite mechanisms compared to the low *η* model (Figure 2E).

### 3.3 Altered Interoception: Mountain-Climbing Task

To investigate the impact of alterations in interoception on balance with regard to minor immediate and major long-term rewards, we used a mountain-climbing task (Figure 3A, B). To assess changes in behavior, three types of measures were used: 1) *S*_*7*_-rate, which is the number of time steps spent in *S*_*7*_ divided by the total number of time steps within a single trial; 2) total-time steps, which is the sum of all time steps within a single trial; and 3) *a*^*00*^-timesteps, which is the total number of times that a small intake (*a*^*00*^) was selected during a single trial. A larger *S*_*7*_-rate (Figure 3C), smaller total-timesteps (Figure 3D) and *a*^*00*^-timesteps (Figure 3E) characterized the priority for long-term, large rewards. The overestimated interoception agent showed a higher *S*_*7*_-rate (Figure 3C), fewer total time steps (Figure 3D), and lower *a*^*00*^-timesteps (Figure 3E). This suggests that agents prioritize reaching the summit and receiving a large reward, even if this means short-term nutritional loss, over immediately receiving a small reward. In the underestimated interoception condition, the HRL models showed no significant changes in *S*_*7*_-rate (Figure 3C) and total-timesteps (Figure 3D), but decreased *a*^*00*^-timesteps (Figure 3E), suggesting *Q*-values of actions that moved away from the end of the episode were larger (Figure 3F; *Q0*-value).

### 3.4 Altered RL Parameters

As mentioned earlier, we endeavored to compare the impact of altering RL parameters, namely inverse temperature (*β*) and discount rate (*γ*), which have been linked to depression, with the effects of modifying interoceptive processing in the HRL model. First, we conducted the intake-after-food-restriction task by manipulating the inverse temperature parameter *β*, which is associated with effects of *Q*-values to the action selections (Schweighofer and Doya, 2003). As a result of manipulating *β* in the HRL model, the reward gained from a single intake increased, while the cumulative punishment also increased significantly (Figure 4A, B). Although this outcome may initially appear counterintuitive, it can be explained as follows. As the optimal behavioral choice in the deficient state (i.e., intake behavior) was reduced, the internal state remained far from the set point (Figure 4E). This, in turn, reduced the proximity to the ideal value of the internal state in the homeostatic space, resulting in greater rewards being obtained from a single intake within the homeostatic space (i.e., at a position where the change in drive per intake was greater; Figure 4B). In fact, when we assessed the total drive within a single episode, it was observed to increase as *β* decreased (i.e., when homeostasis was altered; Figure 4G).

**Figure 4:**
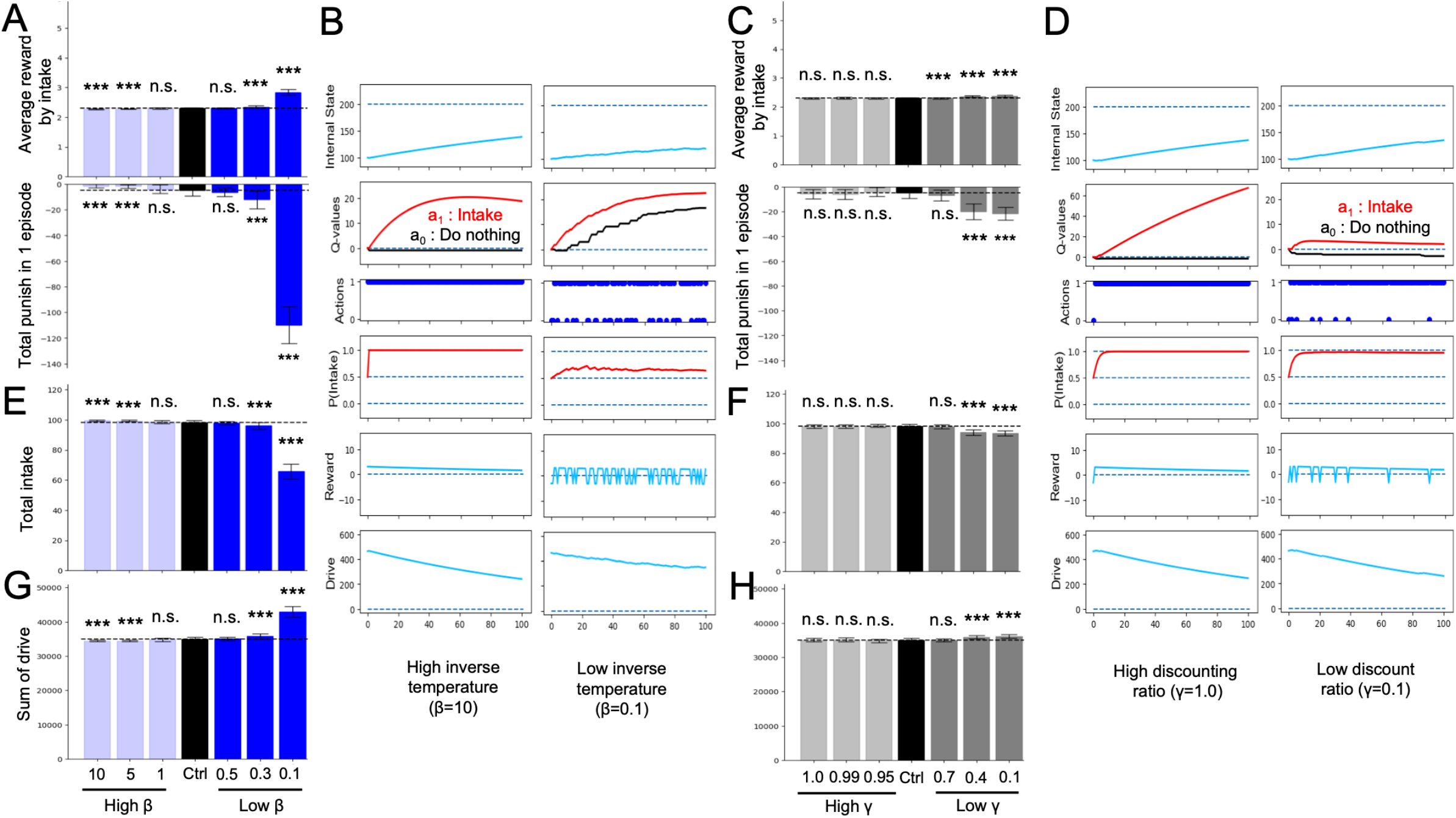
Alterations in reinforcement learning (RL) parameters in the intake-after-food-restriction task. **(A)** Average rewards per intake behavior in single episodes and sum of punishment in each episode determined from the simulations with altered inverse temperature (*β*). **(B)** Example transitions of variables of the altered *β* models. **(C)** Average rewards with all intake behavior in single episodes and sum of punishment in each episode determined from simulations with altered discount ratio (*γ*). **(D)** Example transitions of variables of the altered *γ* models. **(E)** Total number of intakes in single episodes determined from simulated lesion models with altered *β*. **(F)** Total number of intakes in single episodes determined from models with altered *γ*. **(G)** Sum of drive during single episodes of models with altered *β*. **(H)** Sum of drive during single episodes of models with altered *γ*. \*\*\**P* < 0.001 (N = 40); *N*.*S*., not significant. In panels (A), (C), and (E–H), Student’s *t*-test or Welch’s *t*-test was used after Levene’s test.

We also manipulated *γ* in the intake-after-food-restriction task. When *γ* was decreased, the subjects made action selections emphasizing immediate rewards or punishments rather than future ones. This manipulation of the HRL model resulted in a slight increase in the average reward for each intake episode (Figure 4C), increase in the sum of drives in a single episode (Figure 4H), and decrease in total intake (Figure 4F). This is because decreasing *γ* reduced the absolute value of the second term in Eq. 4 (*δ*: TD error), thereby reducing the range of possible values for *δ* and range of possible *Q*-values. Consequently, the differences between *2* actions were reduced as low *β* (Figure 4B, D), impacting the reward and drive similar to that in the low *β* model.

We further examined the effects of changing *γ* and *β* of the HRL model in the mountain-climbing task. Models with decreased discount rate *γ* consistently showed a lower frequency of summit attainment across all metrics compared to the control group (Figure 5A–C). Notably, these models experienced an increase in short-term rewards and stayed at lower altitudes (Figure 5C), underscoring the tendency of smaller *γ* values to prioritize immediate rewards and punishments over distant future rewards. This observation confirmed the simulation’s assumption that models with lower *γ* behave more impulsively, thereby validating the rationality of the mountain-climbing task. In the low *γ* group shown in Figure 5D, the darkest blue line represents the *Q*-value for *a*^*00*^, and the second darkest line indicates the *Q*-value for *a*^*10*^, suggesting that the differences in state-action values reflect the elevation in short-term state-action values. Additionally, increased frequency of stays at states *S*_*0*_ and *S*_*1*_ in this group, as shown in Figure 5D, indicates the decreased climbing performance due to the relative rise in lower state-action values in the low-gamma group. Models with low *β* also demonstrated a reduced frequency of reaching the summit across all measures compared to the control group (Figure 5A, B, C). This can be attributed to the increased randomness in behavior caused by low *β*, leading to less frequent selection of the optimal behaviors necessary for achieving rewards at the summit, especially during periods of nutritional deficiency.

**Figure 5:**
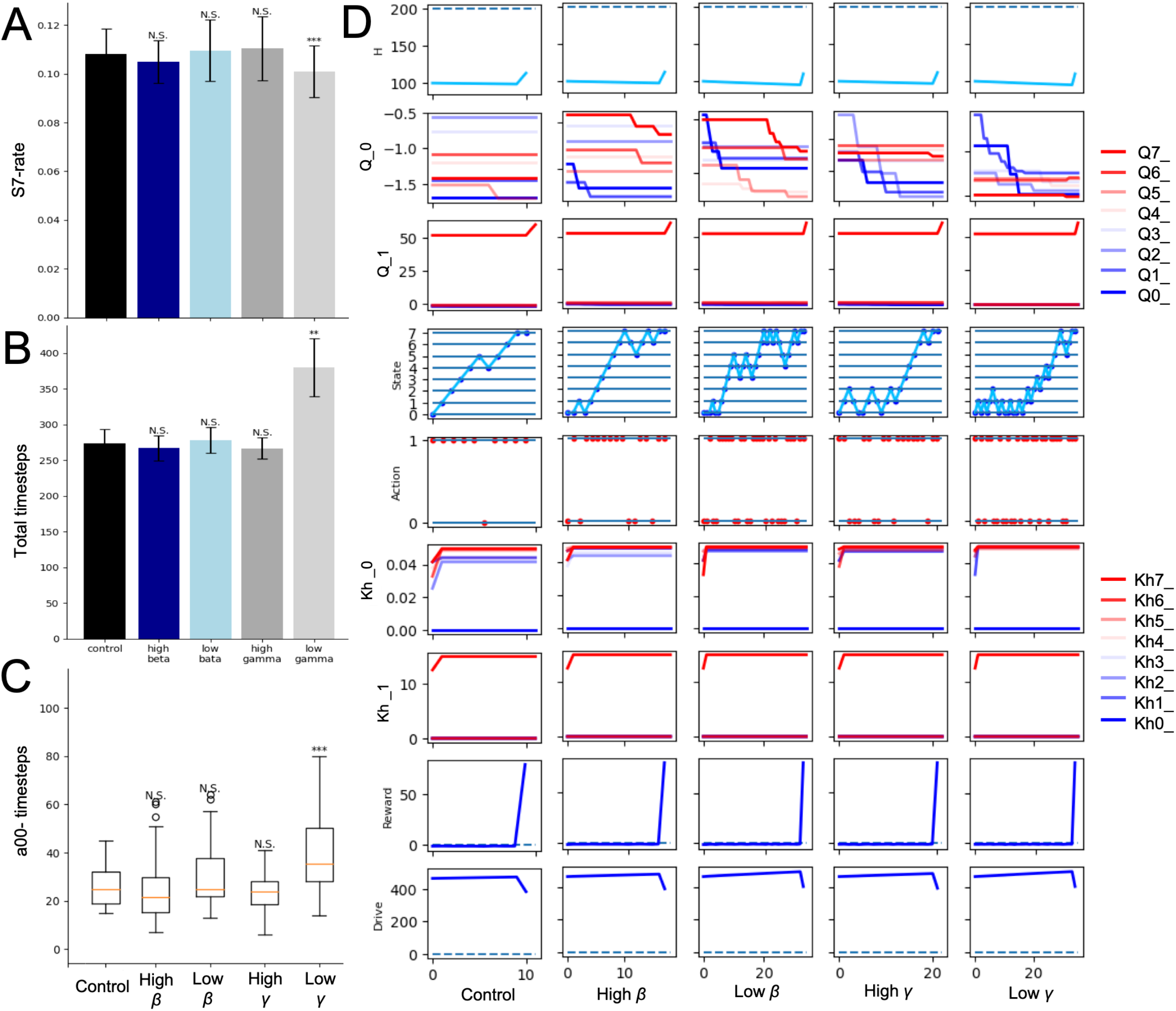
Alterations in RL parameters in the mountain-climbing task. **(A)** Results of the control, low *γ*, and low *β* models referred to the ratios determined from the same calculations as those shown in Figure 3C. **(B)** The same measure of Figure 3D. **(C)** The same measure of Figure 3E. **(D)** Trajectories of each variable in the final episode of low *γ* and low *β* models. *N*.*S*., not significant; \*\**P* < 0.01 and \*\*\**P* < 0.001.

## 4 Discussion

In this study, we attempted to interpret the changes in homeostatic maintenance behaviors and reward system activity by conducting pseudo-manipulation simulations of reduced and exaggerated interoception using the HRL model. Additionally, we compared the effects of modifications in RL parameters associated with depression on decision-making behavior with the effects of changes in interoceptive processing within the HRL model. Through this comparison, we aimed to provide a systems-level explanation of the relationship between phenomena previously considered to be changes in RL parameters, changes in interoceptive processing, and nutrient homeostasis.

In the low *η* model, the difference between the ideal state and the actual internal state was mitigated, resulting in a smaller reward per intake behavior (Figure 2A). This suppressed the learning of the *Q*-value for optimal behavior (intake; Figure 2E). As a result, the agent spent more time in a state in which the internal conditions deviated from the optimal values. Due to this prolonged deviation, the accumulated drive remained high, reflecting the failure of the system to regulate itself effectively (Figure 2C). However, no significant changes were observed during the mountain-climbing task (Figure 3). In the high *η* model, in contrast, the difference between the ideal state and the actual internal state was exaggerated, resulting in a larger reward per intake behavior (Figure 2A). This made it easier to learn the *Q*-value for optimal behavior, and the total drive decreased quickly (Figure 2E). However, this model demonstrated difficulties in acquiring large distant rewards (completing the task) in the mountain-climbing task (Figure 3C, D, E, F). This was due to the tendency to overestimate immediate punishment before a large reward and the small rewards obtained away from the large reward (Figure 3).

The observations in these models were qualitatively similar to depressive symptoms. For example, in the reduced interoception model, we observed a decrease in the range (*R, Q*) of reward system activity (Figure 2A, E). This can be understood as a pattern of reduced activity in the insular cortex, which processes interoceptive information, and in the reward system of patients with depression with reduced appetite. Furthermore, the increase in punishment across tasks (Figure 2B), decrease in optimal intake behavior in the post-dietary intake task (Figure 2D), and increase in total drive (Figure 2C) can be understood as corresponding to depressed patients’ subjective feelings of inadequate internal state maintenance and sustained physical strain. In contrast, in the increased interoception (high *η*) model, increased reward system activity (Figure 2A, E) was observed. This can be understood as a pattern similar to the increased insular cortex activity and increased reward system activity in patients with depression and increased appetite. This increased interoception model demonstrated appropriate behavior in the intake-after-food-restriction task but prevented the completion of the mountain-climbing task. This can be understood to be similar to the deficits in long-term reward-oriented behavior observed in depression. Notably, there were no significant differences in body mass index (BMI) between the groups of subjects who showed clear contrasts in the activities of the insular cortex and reward systems. This is consistent with the fact that the internal state (*H*) did not differ from that of the control model in either the increased or decreased interoception models. Thus, these manipulated interoception models may represent an aspect of the pathophysiology of a subtype of depression characterized by decreased or increased appetite.

In addition, we examined the effects of RL parameters, which are often discussed in relation to depression. Previous behavioral modeling studies of depression using RL models, such as simple Q-learning, have highlighted increased behavioral randomness and low *β* values, tendency to overestimate short-term rewards while underestimating long-term rewards, and association with a reduced *γ* value. Notably, HRL model exhibited trends similar to those of the conventional RL model. In the low *β* model, the tendency for the value of intake behavior increased normally (Figure 4B), but when calculating the probability of action (*P*) from the *Q*-value, the relative magnitude of the two behavioral values was underestimated, resulting in a lower frequency of the appropriate behavior, intake, being selected (Figure 4B). Similarly, the low *γ* model showed the same trend as the conventional RL model. That is, the future value of the behavior is underestimated, and the prediction of immediate reward or punishment strongly influences decision-making. Consequently, the climbing task required more time steps to reach the summit (Figure 5).

Behaviors of these RL parameter modulation models have both similarities and differences with the results of the interoception modulation models. First, low *η*, low *β*, and low *γ* models exhibit similarity in increased drive (Table 1). These results are due to a decrease in the frequency of optimal behavior, resulting from the smaller range of rewards and *Q*-values (low *η* and low *γ*) or difficulty in reflecting the relative magnitude in action probabilities. For the high *η* and low *γ* models, whose performance declined in the mountain-climbing task, responses to immediate rewards and punishments increased, and the reward for each intake was large. However, in the intake-after-food-restriction task, the drive increased in the low *γ* model but decreased in the high *η* model, where high *η* was more adaptive. Although high *η*, low *β*, and low *γ* showed similar increased reward per action, two different mechanisms were involved: high *η* overestimated the reward for a change in internal state of a certain magnitude, whereas low *β* and low *γ* did not alter the evaluation of rewards for internal states compared to the control. However, in low *β* and low *γ* models, more time was spent in regions where the internal state had significantly deviated, and the punishment for failing to choose intake was high.

**Table 1:** Summarized results.

|  | Reward | Drive | Mountain climb |
| --- | --- | --- | --- |
| Low $\eta$ | ↓ | ↑ | → |
| High $\eta$ | ↑ | ↓ | ↓ |
| Low $\beta$ | ↑ | ↑ | → |
| Low $\gamma$ | ↑ | ↑ | ↓ |

These results indicate that individuals showing similar results in one task may have different underlying mechanisms and exhibit different behaviors in another task. Therefore, influence of homeostasis and interoception should be considered when discussing the relationship between RL and depressive symptoms related to nutrition. Although the current model focuses on the deterministic modulation of interoception, actual eating behavior is possibly influenced by various factors, such as hormones and visual stimuli, over different time scales. Therefore, further detailed computational modeling is warranted to understand the physiological homeostasis and mechanisms in psychiatric disorders.

## 6 Author Contributions

YU conceived the study, performed the experiments, and analyzed the data. YU and YY designed the experiments, analyzed the data, and wrote the manuscript. TH and MH critically reviewed and revised the manuscript. All authors contributed to the article and approved the submitted version.

## 8 Acknowledgments

We would like to thank Editage for English language editing, and GPT-4o model for providing valuable language assistance in the preparation of this manuscript.

## 10 Supplementary Material

Supplementary Tables 1 and 2.

## 12 Data Availability Statement

The codes used in this study can be found at: https://github.com/YuukiUchida/Uchida2024DepNutrHRL_prep.

