## SupplementaryTables for "Heterogeneous Appetite Patterns in Depression: Computational Modeling of Nutritional Interoception, Reward Processing, and Decision-Making": SupplementaryTable1.docx

Supplementary Table 1; Intake-after-food-restriction task

| Free parameter | Value | Explanation |
| --- | --- | --- |
| $\alpha^{Q}$ | 0.3 | Learning rate of state-action values |
| $\beta$ | 0.6 | Inverse temperature of action selections |
| $\gamma$ | 0.9 | Discount rate |
| $m$ | 3 | Free parameter of the drive function |
| $n$ | 4 | Free parameter of the drive function |
| $\tau$ | 200 | Attenuation rate of the internal state |
| $\alpha^{\hat{K}}$ | 0.3 | Learning rate of predicted value of increases in the internal state |
| $K$ | 2 | Volume of small intake |
| $H^{*}$ | 200 | The ideal internal state |
| $H_{0}$ | 100 | The depleted initial internal state |
